# Restoration of mitochondrial function to metabolically stressed astrocytes by fusion with mesenchymal stromal cells

**DOI:** 10.1101/2025.02.03.635611

**Authors:** Remy T. Schneider, Matthew R. Rowe, Rebecca H. Dawson, Katerina Komrskova, Jiri Neuzil, Michael V. Berridge, Patries M. Herst, Melanie J. McConnell

**Affiliations:** School of Biological Sciences, Te Herenga Waka Victoria University of Wellington, New Zealand; Malaghan Institute of Medical Research, Wellington, New Zealand; Laboratory of Reproductive Biology, Institute of Biotechnology of the Czech Academy of Sciences, Czech Republic; Department of Zoology, Faculty of Science, Charles University, Prague, Czech Republic; Laboratory of Molecular Therapy, Institute of Biotechnology of the Czech Academy of Sciences, Prague-West, Czech Republic; School of Pharmacy and Medical Science, Griffith University, Southport, QLD, Australia; Department of Radiation Therapy, University of Otago, Wellington, New Zealand

**Keywords:** mitochondria, astrocyte, cell fusion, mesenchymal stromal cell

## Abstract

Mitochondrial networks in eukaryotic cells are dynamic, undergoing fusion, fission, biogenesis and transfer between cells to manage cellular requirements for energy production and biosynthetic components. Mitochondrial transfer has been observed in response to cellular injury and can happen by a variety of mechanisms. To examine whether mitochondria could be restored to injured astrocytes, we established a model of discrete astrocyte-specific, mitochondrial-specific injury in a mixed culture model. Immortalized ρ^0^ astrocytes were generated, then put under lethal metabolic stress by withdrawal of pyruvate and uridine. When cells stressed in this way were grown in the presence of mesenchymal stromal cells, enhanced survival of the ρ^0^ cells was seen, accompanied by the rapid acquisition of an entire mitochondrial networks. Gap-fill ligation and rolling circle amplification for cell-specific mitochondrial genome SNP confirmed that the mitochondrial networks were derived from the mesenchymal stromal donor cells, and both mitochondrial DNA and mitochondrial function were restored to the p0 cells long-term. The presence of two nuclei in the ρ^0^ cells in the early stages of mitochondrial transfer and recovery was observed. This demonstrated that in this instance, the mechanism of mitochondrial rescue was fusion of the ρ^0^ astrocyte and the mesenchymal stromal cell. In support of this, other neural cells were similarly shown to fuse with stressed ρ^0^ astrocytes, suggesting that whole organelles may be donated to restore critical mitochondrial function.

## Introduction

Eukaryotic cells rely on healthy mitochondria for their survival. In addition to being a hub for several anabolic pathways, mitochondria contribute substantially to the cell’s energy budget^1^. The mitochondrial electron transport chain sets up a proton gradient across the inner mitochondrial membrane, which is used by ATP synthase to generate ATP via oxidative phosphorylation. In addition, the electron transport chain is used to support the activity of dihydroorotate dehydrogenase (DHODH), an enzyme essential for the *de novo* synthesis of pyrimidines required for DNA replication and cell division^2^.

Mitochondrial fusion, fission and biogenesis creates a dynamic network that can reshape and resize to satisfy the energy demands of the cell. This dynamism extends to the ability of mitochondria to move between cells. While originally an unexpected finding^3,4^, this trafficking of mitochondria between cells is consistent with both the symbiont origin of mitochondria, and their integral role in maintaining cell viability and cell division.

Mitochondrial transfer has often, though certainly not exclusively, been observed in response to cellular injury. These injuries are often broad acting and can result in damage to other components of the cell as well as to mitochondria. Transfer of mitochondria has been correlated with rescue of the injury in a variety of cell types *in vitro* and *in vivo*^5–7^. Mitochondrial-related rescue from injury could occur via the uptake of soluble small metabolites; mitochondria-containing vesicles docked on the outer leaflet of recipient cell’s membrane; or true inter-cellular mitochondrial transfer^8^. This transfer of mitochondria into the cytosol of another cell has most commonly been observed to occur by the formation of ‘tunneling nanotubes’ between cells, and by generation and uptake of extracellular vesicles containing mitochondria, although other modes of transfer are also possible.

Mitochondrial rescue, by transfer or by other means, may be a critical component in the response to injury. Understanding the cellular response to injury is particularly critical in the nervous system. The brain uses 20% of the total amount of oxygen consumed by the body, with much of that oxygen used to generate energy through oxidative phosphorylation. Neural cells such as neurons and astrocytes are vulnerable to diminished levels of oxidative phosphorylation, as well as loss of the biosynthetic intermediates produced by the TCA cycle. In addition, neural cells are not highly proliferative, meaning that damaged and dead cells are not readily replaced. The cumulative effects of compromised mitochondrial metabolism can be seen in neurodegenerative disorders, where increasing levels of respiratory and metabolic defects are commonly observed as degeneration progresses^9,10^. The astrocyte is increasingly recognized as a critical cell in the function of the brain in health and disease. Indeed, specific mitochondrial injuries to astrocytes are observed in neurodegenerative conditions such as Alzheimers, Parkinson’s and Huntington’s diseases^11–13^.

To examine how astrocytes would respond to mitochondrial injury, we established a model of discrete astrocyte-specific, mitochondrial-specific injury in a mixed culture model. Central to this, a primary murine astrocyte cell line was established, and a metabolically vulnerable ρ^0^ derivative of this cell line was established, by removal of the mitochondrial genome^14^. Because the mitochondrial DNA encodes 13 subunits of the mitochondrial electron transport chain (7 of respiratory complex I, 1 of respiratory complex III, 3 of respiratory complex IV and 2 of respiratory complex V), electron transport and oxidative phosphorylation are lost, so ρ^0^ cells are exclusively glycolytic. This requires supplementation with pyruvate and uridine^14^. Pyruvate is a critical metabolite that can be oxidized in the mitochondria to acetyl coenzyme A and enter the citric acid cycle. In addition, pyruvate can be reduced to lactate in the cytoplasm to support continued glycolysis by re-oxidising the intermediate electron acceptor, NADH to NAD^+^. In ρ^0^ cells in the presence of glucose, the majority of pyruvate is converted to lactate to maintain glycolytic ATP production^15^ This leaves a deficit of pyruvate for conversion to acetyl CoA, TCA cycle activity and thus biosynthetic precursors. The addition of pyruvate in the medium of ρ^0^ cells ensures enough pyruvate is available for both continued glycolysis and TCA cycle activity. Uridine is required to circumvent the block on pyrimidine synthesis. *De novo* pyrimidine synthesis relies on functional electron transport in the mitochondria. Dihydroorotate dehydrogenase (DHODH), the 4^th^ enzyme in the pyrimidine pathway, reduces Coenzyme Q10, located prior to respiratory complex III, in order to oxidise dihydroorotate to orotate.^16^ Mitochondrial competent cells undergoing oxidative phosphorylation continually re-oxidise CoQ10 as part of electron transport. Without electron transport, ρ^0^ cells are incapable of producing pyrimidines, DNA and thus undergoing cell division.^17^ Not supplementing ρ^0^ cells with pyruvate and uridine leads to cell death *in vitro*. ρ^0^ cells *in vivo* obtain OXPHOS competent mitochondria from their surrounding microenvironment.^4^

## 2. Materials and methods

### 2.1 WT and ρ^0^ astrocyte generation

#### 2.1.1 Isolation and immortalization of eGFP-expressing astrocytes

eGFP mice were used to establish a culture of astrocytes, according to published methods.^18^ All experiments using tissue from mice were conducted in accordance with the New Zealand Animal Welfare Act 1999 (www.legislation.govt.nz/act/public/1999/0142/latest/DLM49664.html) and were approved by the Victoria University of Wellington Animal Ethics Committee. Briefly, the cortex and hippocampus were isolated from 5 newborn mouse pup brains, a single cell suspension rendered and all the cells plated in one T75 flask. The flask was incubated at 37°C, 5% CO_2_, for approximately 4 days, until astrocytes were adhered and growing under tissue debris. The cells were then rinsed 3 times with astrocyte media (DMEM high glucose, 100U/mL penicillin-streptomycin, 1 mM sodium pyruvate, 18% FBS) and cultured for an additional 7 days, changing the media every 2 days. When the cells reached over 80% confluence, microglia, oligodendrocyte progenitor cells (OPCs) and neurons, could be identified growing on top of an astrocyte bed. The culture was shaken at 1000 RPM for 2 hours, the microglia-containing supernatant was then discarded, fresh media added, and the culture set to shake, overnight, removing the OPC into the supernatant, which was then discarded, making the culture over 90% astrocytes. The cells were then lifted by incubation with 0.05% Trypsin-EDTA for 5 to 7 minutes, and the astrocytes were seeded at 1×10^6^ cells in a T75 flask. After 7 days, the culture flask reached 90% confluency. Cells were then passaged by the 3T3 protocol (1:3 every 3 days regardless of confluency). The growth of the culture slowed after passage 3, cells exhibited increase in cell surface area with a planar phenotype. At passage 18 the eGFP astrocytes spontaneously immortalized, increasing their growth rate and becoming more compact.

#### 2.1.2 Generation of ρ^0^ eGFP astrocytes

To create ρ^0^ eGFP astrocytes, the cells were treated with 1µg/mL ethidium bromide (EtBr) for 8 weeks in the presence of 1mM sodium pyruvate and 1mM uridine. At this low dose, EtBr preferentially intercalates into mitochondrial DNA over nuclear, where it blocks replication of the mitochondrial genome, causing a depletion of mtDNA over time^14^.

#### 2.1.3 DNA extraction and PCR

Genomic and mitochondrial DNA was extracted from wildtype and ρ^0^ astrocytes using the Zymo Research Quick-gDNA™ MiniPrep (Ngaio Diagnostics, New Zealand). PCR analysis was then used to amplify a 114bp fragment of the *Cytb* gene, encoded in the mitochondrial genome with forward primer (5’-TCC TTC ATG TCG GAC GAG GCT-3’) and reverse primer (5’-ACG ATT GCT AGG GCC GCG AAT-3’). A 148 bp fragment of the ApoB was similarly amplified from the nuclear genome, with forward primer (5’ TCACCAGTCATTTCTGCCTTTG 3’) and reverse ((5’ CACGTGGGCT CCAGCATT 3’). PCR was carried out with the following thermocycling conditions: 2 min at 95°C; 35 cycles of 20 s at 95°C, 15 s at 60°C, 30 s at 72°C; and 2 min at 72°C. PCR products were then analyzed by electrophoresis in a 2% agarose gel, and visualized by ethidium bromide staining.

### 2.2 Cell imaging

#### 2.2.1 Coverslip preparation

Glass coverslips were immersed in 70% nitric acid for 48 hours to etch the surface and facilitate cell binding, then washed 3 times with ddH2O, one wash overnight. The coverslips were then transferred to 100% EtOH, flame-sterilized and placed in 12-well plates. Coverslips were coated with 50 µg/mL poly-D-lysine hydrobromide (PDL) in sodium borate buffer and incubated for a minimum of 2 hours at 37°C. Immediately before use, the coverslips were washed 3 times with filtered ddH2O, the last wash left to sit for at least an hour before seeding cells. Alternatively, coverslips were coated with HistoGrip™ (HG) (ThermoFisher, Auckland NZ) as follows. A working solution was prepared at a 1:50 dilution in acetone. Coverslips were incubated for 2-5 minutes in the HG working solution while on an orbital shaker. The treated coverslips were washed 2x with acetone alone, once with 100% ethanol and subsequently stored in 100% ethanol until used. Cells were seeded onto coverslips for live and fixed imaging.

#### 2.2.3 Mitochondrial DNA staining

All visualization of mtDNA by fluorescence microscopy was performed by staining live cells with 6 µg/mL EtBr for 20 minutes in the appropriate cell culture media. EtBr signal was excited at 300 nm and emission detected at 590nm on the Olympus FV1200 laser scanning confocal microscope (LSCM).

#### 2.2.2 Mitochondrial labelling

Mitochondria were labelled with a 1 mM stock solution of MitoTracker® Red CMXRos (MTRX) (ThermoFisher, Auckland, NZ, prepared by dissolving the lyophilized powder in DMSO). Live cells were incubated for 30-45 minutes in final concentration of 1 μM MTRX at 37°C, 5% CO_2_ and CMXRos visualised at 579nm excitation/599 nm emission on the FV1200 LSCM.

#### 2.2.3 CellTrace Violet and CellTrace CFSE staining

Cells were resuspended in PBS at a concentration of 1 x 10^6^ cells/mL, and a droplet of sufficient volume to produce a 5 μM solution of either CellTrace™ dye was dispensed into the cap of the tube. Tubes were sealed and inverted directly onto a vortex mixer for 20 s to facilitate rapid diffusion of the dye throughout the cell suspension. The cells were incubated for 20 min at 37 °C and unconjugated dye removed by centrifugation and washing in PBS, then resuspended as appropriate. Cells were either analysed by flow cytometry, or plated onto coverslips for imaging. Imaging was carried out on the FV1200 LSCM, using 390nm excitation/455 nm emission for Cell Trace Violet, or 488nm excitation/520nm emission for CFSE.

#### 2.2.4. Rolling circle amplification of mtDNA single nucleotide polymorphisms by gap fill ligation

To establish *in situ* SNP detection, cells were seeded directly on Superfrost microscope slides at 70% confluence by surface area and fixed in 70% EtOH for 15 min at room temperature. To assist enzymatic access to mtDNA, samples were digested in pre-warmed 1 μg/mL pepsin (2500U/mg) in 0.1M HCL for 45 s at 37 °C, submerged in PBS for 30 s to dilute residual pepsin, then placed under fresh PBS. Samples were dehydrated by 3 mins in each of 70%, 85%, 100% EtOH and allowed to dry. Up to 8 Secure-seal™ 9 mm diameter hybridisation chambers were fitted to dry slides to form reaction wells for all reactions. Through the 1 mm reagent access ports, samples were rehydrated through a 100, 85, 70% EtOH series for 3 min at each step before a final replacement with PBS.

To prepare mtDNA fragments *in situ*, restriction digest and exonucleolysis were performed in a single reaction (0.5 u/μL of DraI, 0.4 u/μl T7 exonuclease and 1 x NEB Cutsmart buffer, 50 μL per well), and incubated for 30 min at 37 °C. After digestion, each chamber was rinsed gently with RCA wash buffer (0.1 M Tris-HCL pH 7.5, 0.15 M NaCl and 0.05% Tween-20). SNP-specific padlock probe (PDP) were hybridised at 0.1 μM, and ligated with 2 μM of each 5’ phosphorylated gap-fill hexamer and 2μM of each blocker hexamer for the desired reaction targets, 0.1 U/μL T4 DNA Ligase in 1 x T4 DNA Ligation buffer (ThermoFisher). The reactions were supplemented with 0.2 μg/μL BSA, 200 nM ATP and 200 mM NaCl in a final volume of 50 μL per reaction. After 30 min incubation at 37°C, each chamber was rinsed with RCA wash buffer. Fresh RCA wash buffer was supplied to each chamber and the samples were incubated for a further 5 min at 37°C. Each chamber was rinsed 3 times with RCA wash buffer to remove unligated PDPs and hexamers. All probe sequences given in Supplementary Table 1.

To perform rolling circle amplification from circularised PDPs, a reaction mixture was added to each port, consisting of 1 U/μL φ29 polymerase (Lucigen) 0.25 mM dNTPs, 0.2 μg/mL BSA, 5% glycerol and 1 x Phi29 reaction buffer (Lucigen) up to a final volume of 50 μL. Reagent access ports were sealed with SecureSeal™ tabs and placed at 37 °C for 3 h to allow amplification of rolling circle products. After brief rinse with RCA wash buffer, 200 nM of each PDP-paired fluorescent detection oligo (in 20% formamide, 2 x SSC solution) was hybridized at 37 °C for 30 mins. Samples were washed briefly 3x times in RCA wash buffer and dehydrated by 3 mins at 70%, 85% and 100% EtOH, fitted with coverslips in VectaShield™ antifade mounting medium (with or without DAPI) and sealed with clear nail polish. Fluorescent signal was detected at the appropriate wavelength on the FV1200 LSCM.

### 2.3 Bone-derived mesenchymal stromal cell cultures

All experiments using tissue from mice were approved by the Victoria University of Wellington Animal Ethics Committee and conducted in accordance with the New Zealand Animal Welfare Act 1999 (www.legislation.govt.nz/act/public/1999/0142/latest/DLM49664.html).

#### 2.3.1. Preparation of bone-derived cells

The femur and tibia of 12 to 16-week-old B6D2 mito-DsRed, or DsRed:Balb/C F1 mice were isolated and the bones cleaned thoroughly of muscle and connective tissue using fine-meshed gauze. After transferring the bones to a clean 10cm dish with 3 mL of astrocyte media, the epiphyses were removed using a scalpel and remaining compact bone cut into ∼1mm fragments. The bone fragments were washed a few times with media, transferred to a T25 flask, one set of leg bones per flask, and digested with 4 mls of Collagenase II (1mg/mL) (Thermo Fisher, Auckland, NZ) for 2hrs at 37°C, shaking at 200rpm. The digestion was sufficient when the bone fragments no longer clumped together. Subsequently, the bones were washed a few times with media, 5 mL fresh media added, and the flask incubated at 37°C, 5% CO_2_, for 3 days to remove the non-adherent hematopoietic cells. The bone fragments were then transferred to a 10mL dish with media and splintered further with a scalpel. Bone fragments of one mouse were distributed across 15 to 18 wells of a 12-well plate, or equivalent. Stromal cells began migration out of the bone at 2 days and expanded, with proliferation and further migration from bone, to a suitable population size for co-culture at 4 days. Bone fragments were then removed from each well by flooding with 2x the media volume, tilting the plate, allowing the bone fragments to collect at the well periphery and removing them with forceps. The supernatant from each well was saved and returned after the bones were removed and half of the media was changed every two days until the cells were used for an experiment.

### 2.4. Flow cytometry

The media from all conditions was removed to a fresh tube and cells were detached by incubation with 0.25% Trypsin-EDTA for 5 minutes at 37°C. After neutralizing the trypsin with FBS-containing cell culture media, the previously collected media was added to the corresponding cell suspension, to ensure retention and quantification of both live and dead cells.

#### 2.4.1 Assessment of GFP expression

Each cell pellet was resuspended in 1x PBS supplemented with 2% FBS. Propidium iodide (BD Biosciences) was added at 5 μg/mL, or DAPI at 1 μg/mL to determine viability by dye exclusion. Cells were analysed a BD Canto II flow cytometer for eGFP fluorescence, which was measured with 488nm excitation and 520nm emission.

#### 2.4.2 Assessment of apoptosis

This was assessed via exposed phosphatidylserine (PS) on the outer leaflet of the plasma membrane, by adding 3 µL APC-labelled Annexin V (BD Pharmingen, 550475) to each 100 µL cell suspension. The cells were incubated with APC Annexin V for 15 minutes in the dark at room temperature. After adding 400 µL binding buffer to dilute the staining solution, 3 µL propidium iodide (PI) (BD biosciences, 556463) was added, and after a 5-minute incubation at room temperature, the stained cells were analyzed using a BD Canto II flow cytometer, with excitation 633nm/emission 660 nm.

#### 2.4.3 Expression of cell surface markers

These were measured using antibodies to cell surface proteins: CD105-APC (REA794), Miltenyi Biotec; CD73-FITC (TY/11.8), Miltenyi Biotec; CD90.2-VioBlue (30-H12), Milteny Biotec; CD45-BV510 (30-F11), BD Biosciences; CD11b-APC-Cy7 (M1/70), BD Biosciences. CB-MStC cells were harvested by trypsinisation, washed in FACS buffer (PBS, 10% FBS, 1% EDTA) and incubated with 250 μg Fc Block (BD Biosciences) for 30 minutes at 37°C, to block non-specific interactions between cells and the conserved Fc domain of the antibodies. Fluorophore conjugated antibodies were used to label cell surface epitopes. Between 1×10^5^ and 1×10^6^ cells were stained in 50μL on ice for 30 minutes, protected from light, before being washed twice in FACS buffer and resuspended in 500μL FACS buffer with 100 ng/mL DAPI. Single antibody stains were prepared on beads and ‘fluorescence minus one’ FMO controls were prepared on untreated cells. These were used to to determine spectral overlap and set appropriate fluorescence and laser levels on the Fortessa flow cytometer. Flow cytometry data was analysed using FlowJo (BD Life Sciences).

## 3. Results

### 3.1 Generation of wild-type and rho0 eGFP astrocytes

To investigate the response to mitochondrial deficit in neural cells, we developed a ρ^0^ eGFP astrocyte cell line. This line is dependent on pyruvate and uridine for growth, and a lethal metabolic or bioenergetic stress can be selectively induced in these cells by depletion of pyruvate and uridine from the media.

First, primary astrocytes were isolated from the cortex and hippocampus of neonatal eGFP C57BL6 transgenic mice, on the basis of adhesion^18^. They were passaged using the 3T3 protocol - split 1:3 every 3 days - until they underwent spontaneous immortalization at p18, characterized by increased proliferation rate and reduced size. The immortalized astrocytes maintained GFP expression, as well as expression of the astrocyte marker GLAST (Supplementary Fig 1).

Once the eGFP astrocyte line was established, mitochondrial DNA was depleted by long term culture with 1 μg/mL ethidium bromide, supplemented with pyruvate and uridine. Low dose ethidium bromide preferentially binds to mitochondrial DNA, preventing mitochondrial genome replication, meaning that mtDNA was lost by dilution as cells divided in EtBr.^14^ After 8 weeks, the loss of mitochondrial DNA in ρ^0^ eGFP was confirmed by PCR for the mitochondrially encoded gene cytochrome b (*Cytb*). No Cytb could be amplified, confirming the ρ^0^ phenotype of the eGFP astrocytes (Figure 1A).

**Figure 1:**
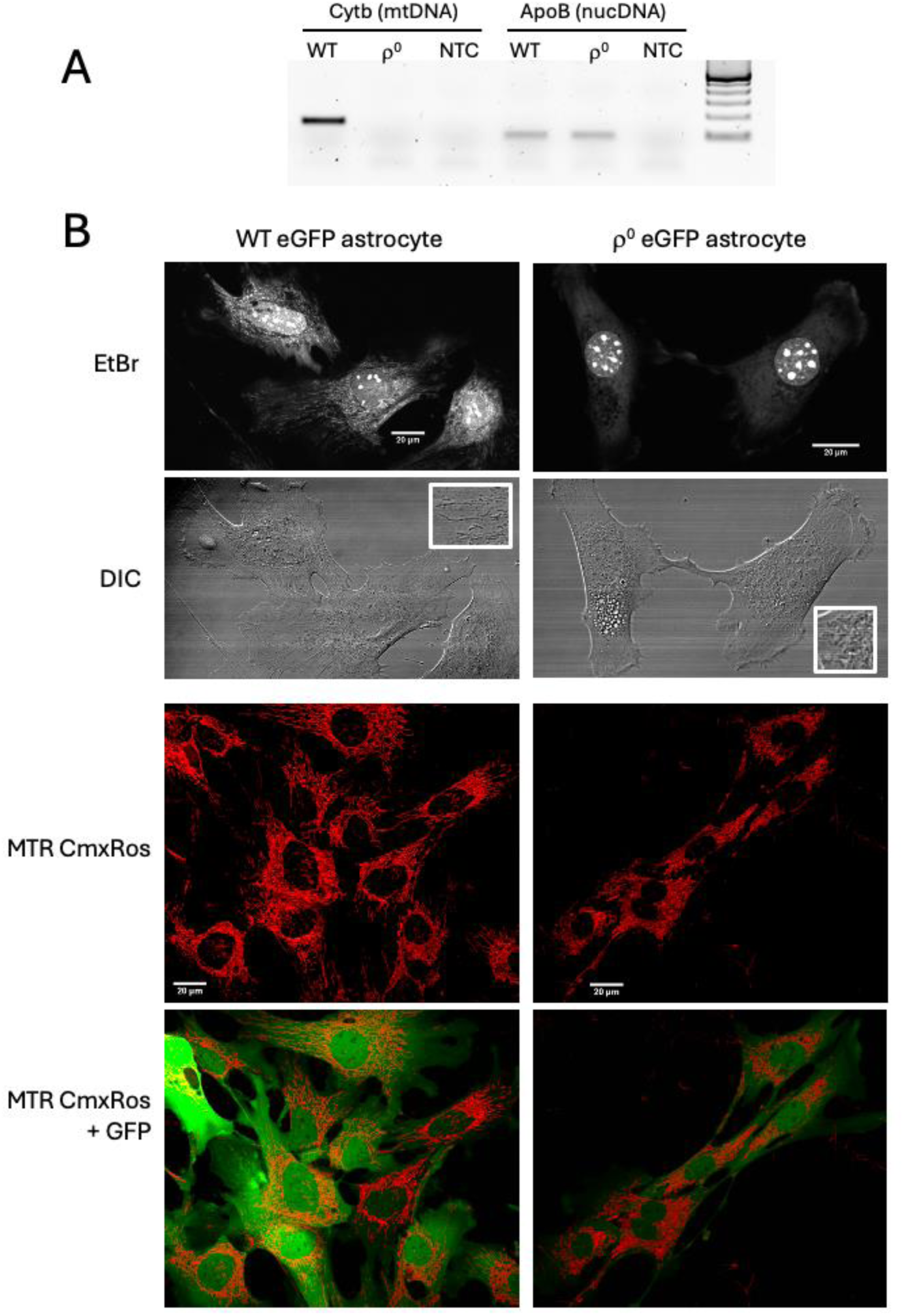
Characterisation of ρ^0^eGFP astrocytes. (A) PCR was carried out for the mitochondrial genome (Cytb, left) and nuclear genome (ApoB, right) from wildtype (WT) and ρ^0^ cells, with water used as the no template control (NTC). (B).Ethidium bromide (EtBr) fluorescence was used to visualise mitochondrial DNA and the nucleolar regions of the nucleus (top), while a DIC image was taken at the same magnification. Scale bar, 20 microns. (C) Mitotracker CmxRos was used to label mitochondria, with eGFP shown in the lower panels. Scale bar, 20 microns.

The loss of mitochondrial DNA was further confirmed by labelling the cells with ethidium bromide. Consistent with prior data^4^ ethidium bromide was incorporated into the mitochondrial networks of the wild-type cells giving a signal throughout the cytosol, but in the ρ^0^ cells only the nucleus could be visualized, in particular the nucleolar regions (Fig 1B). Further, while the untreated cells had long filamentous mitochondrial networks, mitochondria in the ρ^0^ eGFP astrocytes formed a fragmented network, with punctate and swollen mitochondria, as visualized by direct DIC microscopy, and staining with the mitochondrial dye, MTRX. The uptake of MTRX confirmed that ρ^0^ cells maintained a mitochondrial membrane potential in the absence of mitochondrial electron transport (Figure 1B), consistent with published data on ρ^0^ mitochondrial membrane potential, which is maintained through ATPase activity using glycolytic ATP^2^.

Although ρ^0^ eGFP astrocytes had lower eGFP fluorescence and altered morphology compared to the parental WT eGFP astrocytes, viability of the ρ^0^ cells in pyruvate-uridine supplemented growth media was similar to wild-type cells (Supplementary Figure 2).

### 3.2 Injury of p0 eGFP astrocytes by pyruvate uridine deprivation

ρ^0^ cells depend on pyruvate and uridine for survival. When cultured with pyruvate/uridine, the levels of cell death and apoptosis were similar between ρ^0^ and WT eGFP astrocytes, although proliferation rate of the ρ^0^ cells was lower than the WT astrocyte line, as expected from a purely glycolytic cell line. Cells were metabolically ‘injured’ by removal of pyruvate/uridine, and viability monitored for 7 days, using propidium iodide uptake and loss of cytosolic GFP as indicators of compromised membrane permeability, and thus viability. When pyruvate/uridine was removed from the growth medium of ρ^0^ eGFP astrocytes, cell death increased 4-fold from ∼9% at day 2, to ∼36% at day 3, as determined by the GFP^-^/PI^+^ cell population, and further increased from Day 3 to Day 4. By Day 4 only half the number of GFP^+^/PI^-^ cells remained compared with Day 2 (Figure 2A). Removal of pyruvate and uridine separately indicated that pyruvate was the most important factor for survival of the ρ^0^ eGFP astrocyte (Fig 2B).

**Figure 2:**
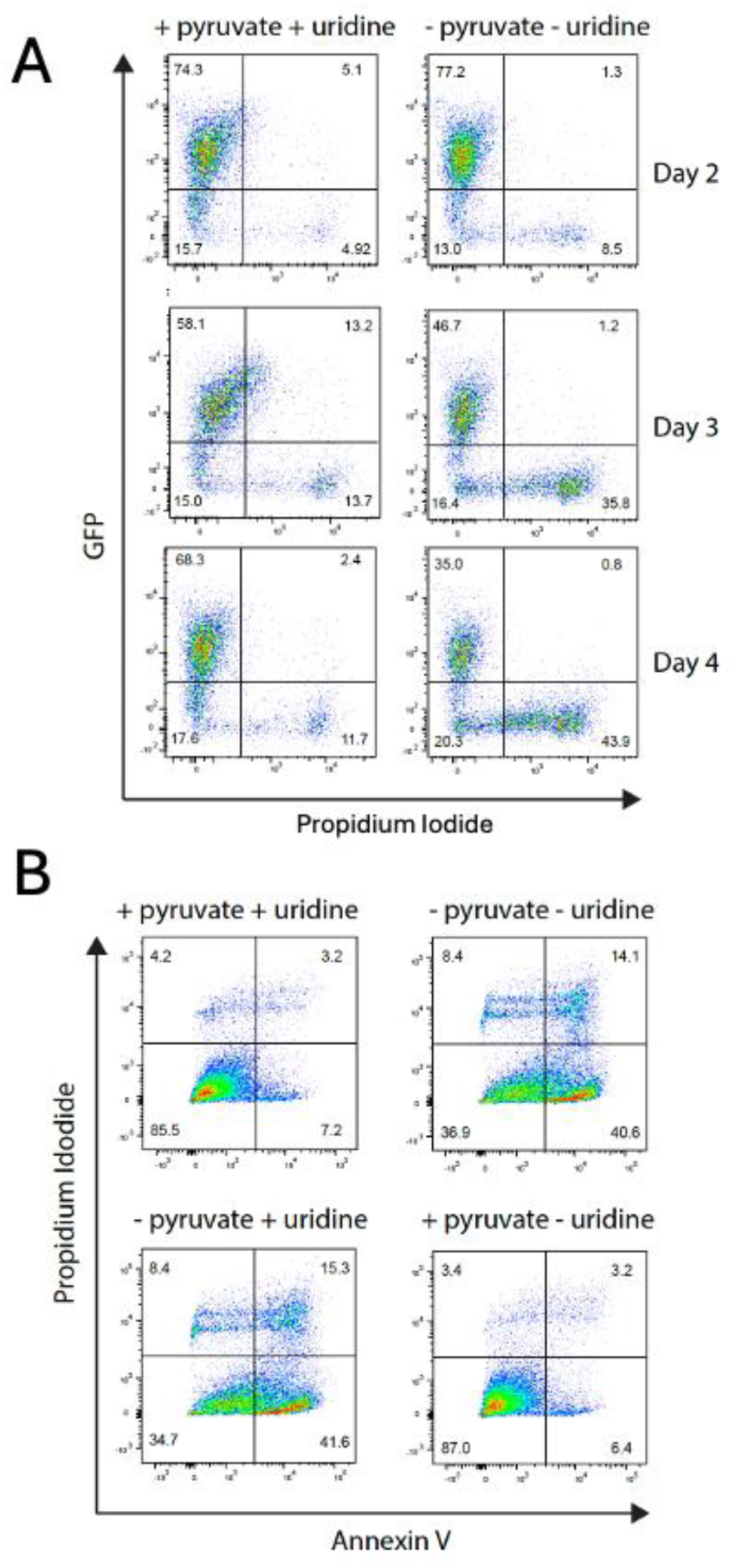
Pyruvate/uridine removal induced death by apoptosis in r^0^ eGFP astrocytes. (A) Propidium iodide update (x-axis) as a measure of dead cells, with consequent eGFP loss (y-axis). (B) At day 4 in the absence of either pyruvate or uridine as indicated, annexin V (x-axis) versus propidium iodide uptake measured early (AnxV+ PI-) and late (AnxV+PI+) apoptosis.

### 3.3 Generation of bone-derived mesenchymal stromal cells

Much of the mitochondrial transfer literature demonstrates that mesenchymal stem and stromal cells (MStC) are effective mitochondrial donors, so we isolated primary mesenchymal stromal cells from compact (a.k.a cortical) bone (CB-MStC).^19^ Bone contains a diverse array of cell types that will migrate out from bone fragments, including osteoblasts, osteoclasts and osteocytes at a range of differentiation stages, hematopoietic progenitors, myeloid cells, mesenchymal stem cells and their stromal descendants.^20^

Mice containing red fluorescent mitochondria were used as the source of stromal cells in our experiments. These animals were generated with a mitochondrial targeting signal (MTS) cloned in frame into pDsRed22-N1 to make transgenic mice that ubiquitously expressed mitochondria-targeted DsRed^21^. The femur and tibia of these mito-DsRed2 mice were flushed of bone marrow, then fragmented and plated directly in glass-bottomed confocal microscopy wells, and cells allowed to migrate out from bone fragments (Fig 3A). After 3 days, suspension cells were removed, leaving multiple adherent cell types with red fluorescent mitochondria (Fig 3B).

**Figure 3:**
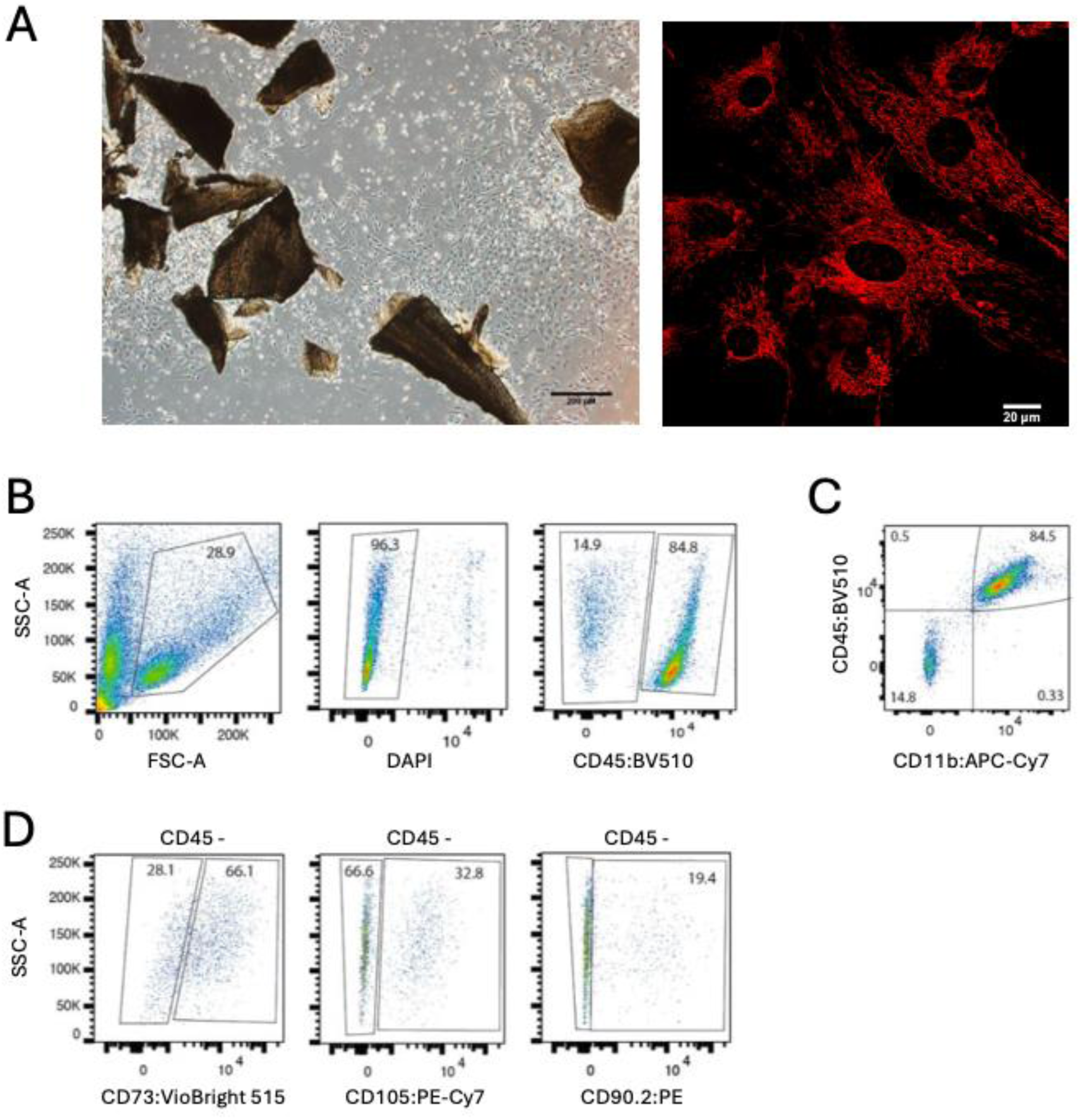
Hematopoietic and mesenchymal stromal cells cultured from compact bone of mitoDsRed mice. (A). Cells migrating from fragments of bone, scale bar 200 micron (left) and mitochondria visualized using MitoDsRed expression, scale bar 20 micron (right). (B) After 1 week in culture, live cells were gated on CD45 (C) CD45+ cells were stained with antibodies against CD11b and CD14. (D) CD45-cells were stained with antibodies against CD73, CD90.2 and CD105. Gates around all populations of cells established on FMO and single-stained controls.

Flow cytometry was used to carry out a preliminary characterization of the major cell types found in the bone-derived stromal cells. The majority (∼67%) of DsRed2 CB-MStC were CD45+ cells, the vast majority of which (98.6%) were CD11b myeloid cells. Most of these (75%) were also CD14+, indicative of macrophage and osteoprogenitor lineages^22^.

The CD45-population (∼33%) included stromal cells that were positive for CD73 (14.9%), CD90.2 (29.5%), and CD105 (35%), indicating the presence of mesenchymal progenitor cells (Figure 3). There were double-positive cells, with 8-10% of the CD45-cells being CD73+CD90+ (Supplementary Fig 3), but very few (<1%) cells that were positive for all three markers, suggesting that while undifferentiated mesenchymal stem cells were largely absent from the cultures, their descendants were present.

### 3.4 Mesenchymal stromal cells only donated mitochondria to ρ^0^ eGFP astrocytes in the absence of pyruvate and uridine

Once the adherent primary mito-DsRed22 CB-MtSC stromal cell colonies reached approximately 500 µm in diameter, ρ^0^eGFP astrocytes were added and both cell types co-cultured in the presence of pyruvate and uridine while the ρ^0^ cells established themselves. After 3 days in co-culture, pyruvate and uridine were removed, then 7 days later, cells were imaged, and compared to co-cultures that retained pyruvate for the full 10 day period (Figure 4).

**Figure 4:**
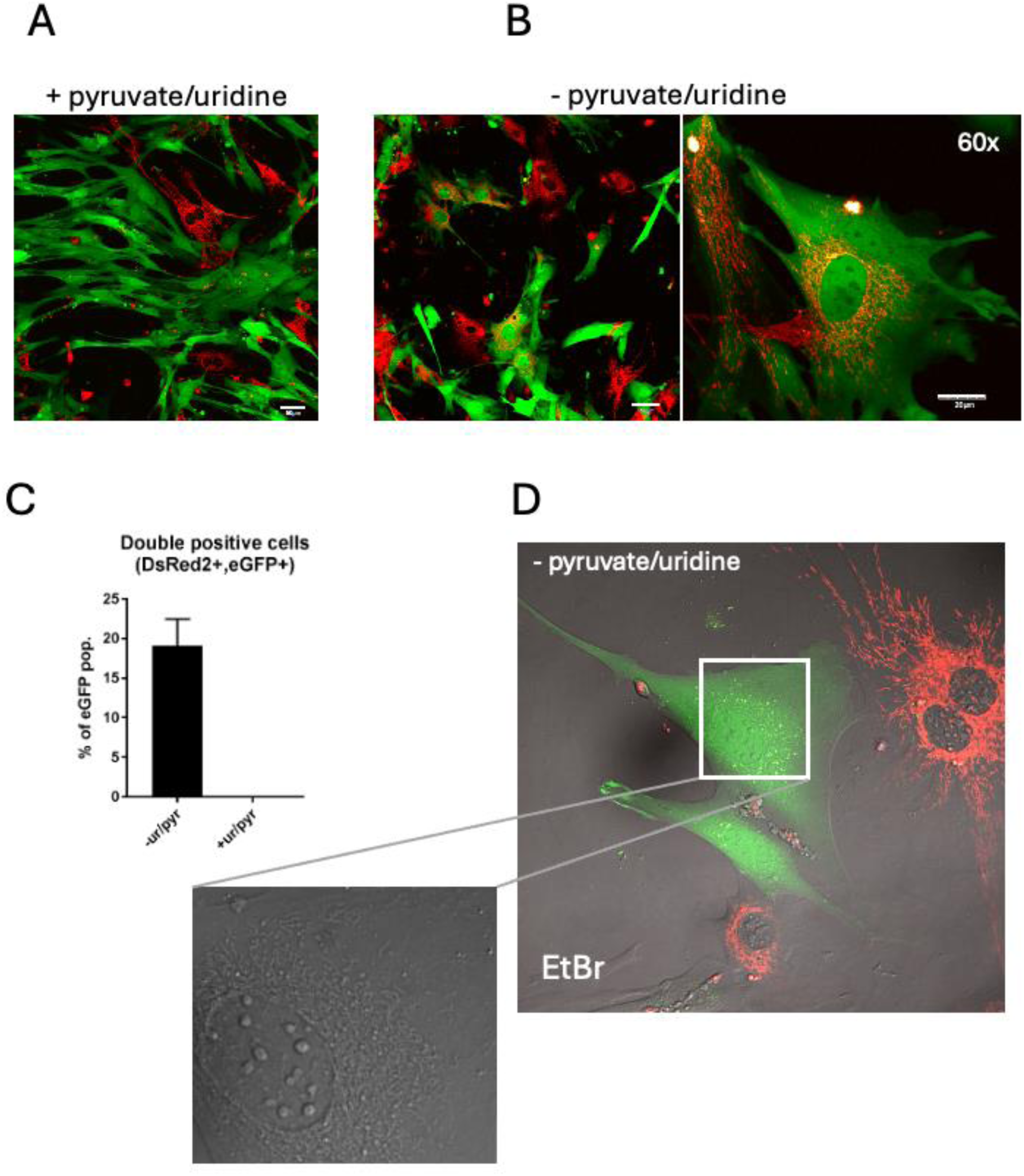
Mitochondrial networks are restored in ρ^0^eGFP astrocytes only after the removal of pyruvate/uridine. Transfer of mitochondrial networks from mitoDsRed mesenchymal stromal cells, mitochondria in red, in co-culture with ρ^0^eGFP astrocytes (green). (A). Co-culture in presence of pyruvate and uridine. Scale bar, 20 micron. (B) Co-culture in the absence of pyruvate and uridine. Left panel, scale bar 20 micron; right panel scale bar 50 micron. (C) The number of eGFP+ cells with red mitochondria was determined from >10 fields, in the presence and absence of pyruvate and uridine. Error bar, standard deviation. (D) Ethidium bromide labelling of mitochondrial DNA in the transferred mitochondrial network, and the perinuclear distribution of the network.

In ρ^0^eGFP astrocytes and mito-DsRed2 CB-MStC co-cultured in the presence of pyruvate/uridine, the red fluorescent signal was limited to the primary mito-DsRed cells, with occasional cell-associated mitochondrial particles visible on and between ρ^0^eGFP astrocyte membranes (Fig 4A). The DsRed mitochondrial particles remained on the cell surface and were never incorporated into the GFP+ve cells. However, in co-cultures deprived of pyruvate and uridine, many ρ^0^ eGFP cells remained viable for 7 days and longer, indicating they had derived support from secreted factors in the media, supplied by the mitoDsRed2 CB-MStC. In addition, almost 20% of the ρ^0^ eGFP cells were repopulated with DsRed mitochondria (Figure 4B, C), forming an extensive mitochondrial network. The presence of mitochondrial DNA in these networks was confirmed by ethidium bromide staining, and the typical perinuclear distribution of the network was observed (Fig 4D).

During an extended period of time (60-90 days) in co-culture, the primary CB-MStC died, having rescued the former ρ^0^ cells by transplant of functional mitochondrial networks, able to thrive in the absence of pyruvate and uridine. By this stage, the mitochondria were no longer fluorescent - the DsRed transgene was expressed from the nuclear genome of the CB-MStC cells, and did not transplant with the mitochondria – but the mitochondrial networks were stable and mitochondrial DNA was routinely detected thereafter.

### 3.5 Mitochondrial genotyping confirmed that mesenchymal stromal cells were the source of mitochondria

It was possible, although very unlikely, that the mitochondria of ρ^0^ eGFP cells could have contained very low levels of mtDNA resulting from incomplete depletion during the initial ethidium bromide treatment. To formally confirm that the CB-MStC was the source of functional mitochondrial networks in ρ⁰ cells, mitochondrial genotyping was used. Male DsRed C57B/6J mice were crossed with female BALB/cByJ, to generate an F1 mouse with DsRed expression and a BALB/cByJ mitochondrial genotype. While the mitochondrial genome is highly conserved among laboratory mice strains, sequencing of the tRNA-Arg locus in the DsRed:F1 mice and the parental eGFP astrocytes confirmed that there were single nucleotide polymorphisms, at positions 9348 and 9461 of the mitochondrial genome, that differentiated the two strains of mice. The eGFP astrocytes maintained the C57BL-6 mitochondrial genotype of 9348-G and 9461-T, while the DsRed2:F1 mice had the Balb/C genotype 9348-C and 9461-A, courtesy of their maternally inherited mitochondria.

This difference was used to determine the origin of mitochondrial DNA within the restored networks of ρ^0^ eGFP astrocytes. An *in situ* genotyping assay^23^ was developed using rolling circle amplification (RCA) with a SNP-specific oligonucleotide padlock probe for each genotype (Fig 4A). The astrocytes in the co-culture were stained with an antibody for the astrocyte-specific glutamate transporter EEAT2/GLAST to positively identify them^24^, as eGFP fluorescence was lost during the RCA protocol. Importantly, successful RCA products in the ρ^0^ eGFP astrocytes were only generated from the probe specific for SNP9461-A, the Balb/C genotype. This indicated that no C57Bl/6 mitochondrial DNA was present in rescued ρ⁰ cells, and that mtDNA in the recovered ρ^0^ eGFP astrocytes had indeed originated from DsRed:F1 stromal cells. Further, confocal microscopy through the XZ axis confirmed that the rolling circle products were inside the cell, demonstrating that the mitochondrial DNA embedded within the body of the cell was the target of the RCA (Figure 5D).

**Figure 5:**
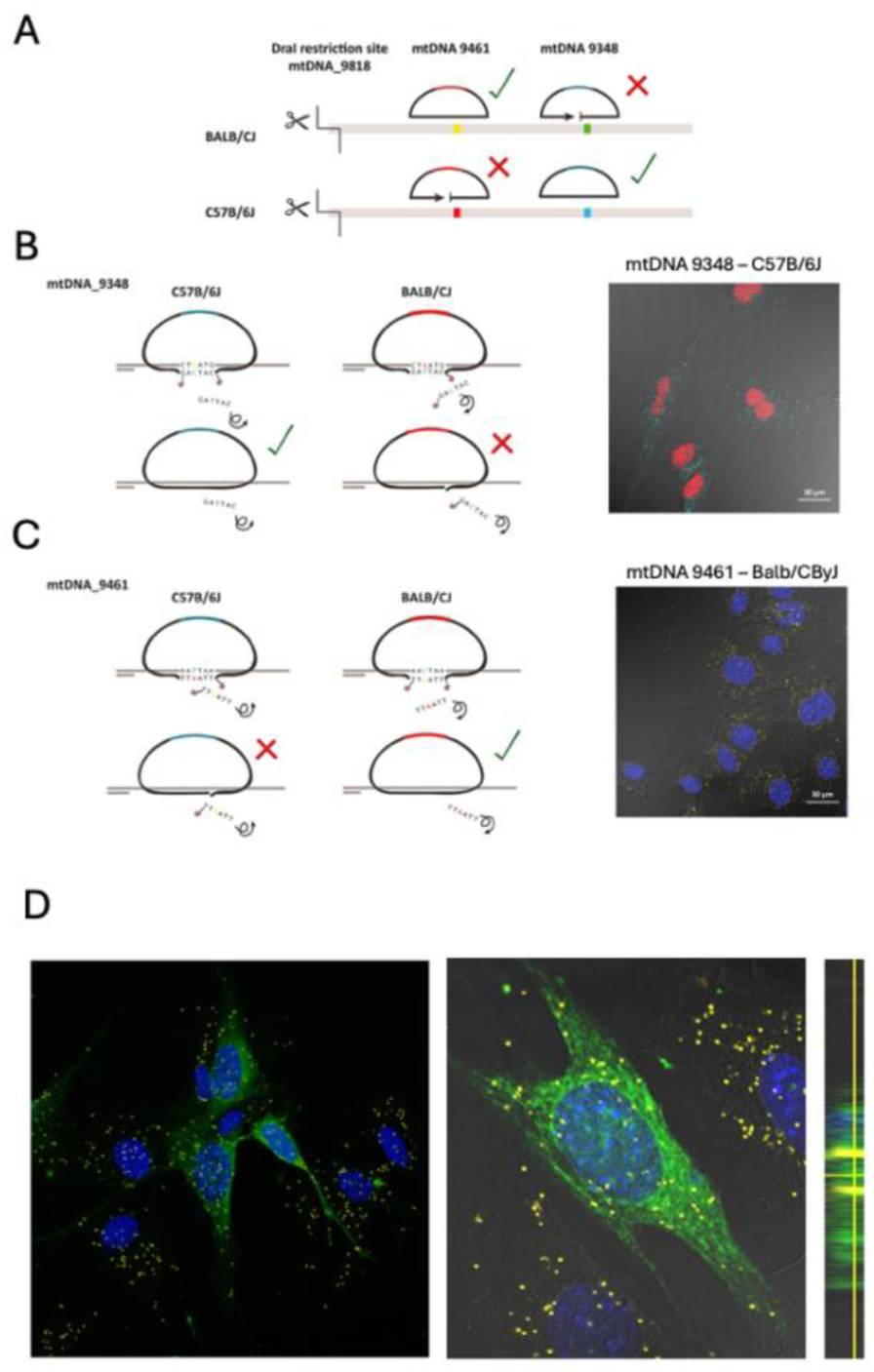
Mitochondrial genotyping by rolling circle amplification (RCA) confirms transfer of mitochondria between cells. (A). Schematic of the probes specific for the C57B/6J mitochondrial SNP 9348-G, and the BALB/C mitochondrial SNP 9461-A. The mitochondrial genome was digested with DraI, which cut ∼400 bases from the two SNPs, at base 9818. Exonuclease makes the DNA single-stranded from the DraI site and beyond the SNPs to allow the probes and padlock oligonucleotides to bind. (B) Detection of the C57B/6 mitochondrial genotype in wild-type C57B/6 eGFP astrocytes cells. RCA was carried out with the 9348-T oligonucleotide padlock, which binds to the A allele in the C57B/6 genome, and allows the Phi polymerase to complete the rolling circle. The amplified sequence was detected with a fluorescent probe, shown in cyan, using confocal microscopy. (C). Detection of the BALB/C mitochondrial genotype in BALB/C bone marrow derived cells. RCA was carried out with the 9461-T oligonucleotide padlock, which binds to the A allele in the BALB/C genome, and allows the Phi polymerase to complete the rolling circle. The amplified sequence was detected with a fluorescent probe, shown in yellow, using confocal microscopy. (D) BALB/C specific 9461-A RCA was carried out in the ρ0eGFP astrocyte + CB-MStC co-cultures. All cells were stained with an antibody against the astrocyte-specific glutamate transporter EAAT2, labelled with a green fluorophore. Left and centre panels are X-Y projections of the confocal image, while the small panel on the right is a X-Z projection.

### 3.6 Mitochondria rescue of stressed ρ^0^ eGFP astrocytes occurred by fusion with stromal cells

It seemed unlikely that a limited number of mitochondria had arrived in each cell via tunneling nanotube^25,26^ or extracellular vesicle^27,28^, and then reconstituted an entire branched network, over less than a week. Instead, we hypothesized that cell fusion occurred between the energetically and biosynthetically stressed ρ^0^ eGFP and the primary mito-DsRed CB-MStC. In support of this, double positive eGFP+ DsRed+ cells were often seen to have more than one nucleus (eg Fig 4B).

To examine multi-nucleate cells, co-cultures were imaged at several points following pyruvate/ uridine withdrawal (Fig 6). Representative images of DsRed2^+^ eGFP^+^ cells at various days after pyruvate/uridine removal showed cells at early timepoints (days 1 and 3) that were large, with a reticulated, tubular mitochondrial network and multiple nuclei with well-defined borders (Fig 6B). The fusion was an early event following the loss of pyruvate and uridine - in the first 24 hours of pyruvate/uridine withdrawal, there were on average 2.5 nuclei per double-positive eGFP+DsRed+ cell. This peaked at ∼3 nuclei per eGFP+DsRed+ cell at 4 days. Beyond this point, the number of nuclei per cell dropped, with most, but not all, double positive cells becoming mono-nucleated from day 6 onwards. No multi-nucleate ρ^0^ eGFP cells were observed in the plus pyruvate/uridine condition. This confirmed that fusion had occurred, and that it was specific to astrocytes undergoing extreme bioenergetic and biosynthetic distress.

**Figure 6:**
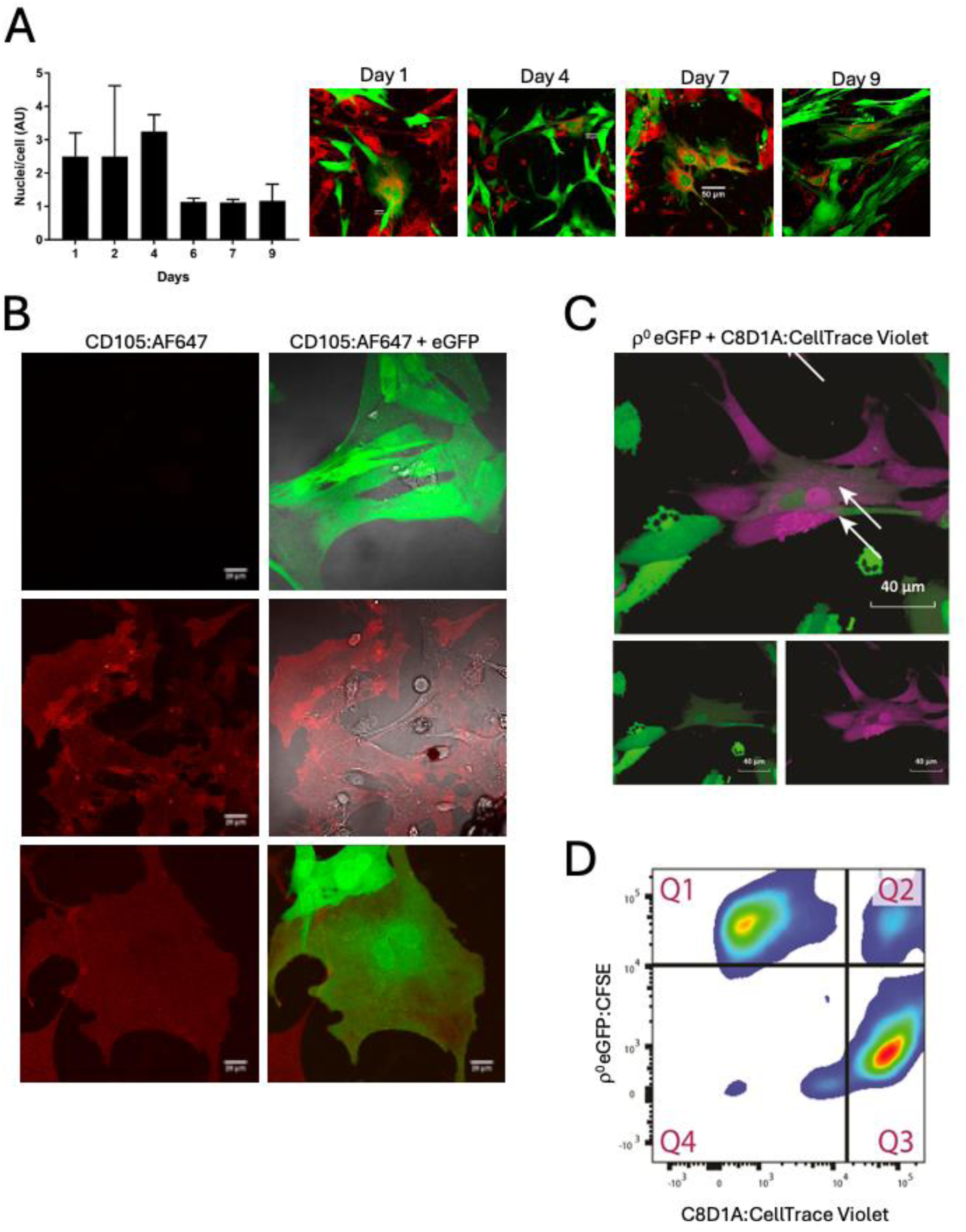
Cell fusion as a method of mitochondrial restoration. (A) The number of nuclei per cell was counted in >50 cells from 3 replicate co-cultures at different timepoints after the removal of pyruvate/uridine, from day 1 to day 9. Error bar, standard deviation. Representative example images are shown on the right, scale bar (50 micron) applies to each image. (B) Co-staining for CD105+ (red) and eGFP (green) in ρ0eGFP astrocyte + CB-MStC co-culture, image acquired by confocal microscopy. Scale bar, 20 micron. (C) Co-culture of ρ0eGFP astrocytes with the C8D1A astrocyte cell line stained with CellTrace Violet (CTV). White arrows indicate dual-labelled cells with both green and violet staining and dual nuclei. (D) Double-positive fused cells were quantified by flow cytometry. Cells in Q1 are GFP+, in Q3 are CTV+ and cells in Q2 are positive for both. Data representative of 3 independent replicate experiments.

To begin to determine which cell type from compact bone had fused with the ρ^0^ eGFP astrocytes, co-cultures were stained with either CD45 for hematopoietic cells, or CD105, the most abundant of the mesenchymal stromal cell markers. There were no cells observed that expressed both eGFP and CD45 - while CD45+ cells were frequently closely associated with the eGFP cells, they remained distinct (data not shown). However, cells homogenously positive for both eGFP and CD105 were identified, indicating that fusion of a ρ^0^ astrocyte and a mesenchymal cell had occurred (Fig 6B). Fused cells also had 2 distinct nuclei, while surrounding cells expressed either CD105 or eGFP, and were mono-nuclear.

The CB-MStC used as a mitochondrial donor in these experiments included mesenchymal lineage cells. However, in our experiments these CD105 cells were derived from bone, which has little immediate relevance to the neural microenvironment. We tested whether a brain-derived cell could similarly act as a mitochondrial donor. The murine astrocyte C8D1A cell line was stained with CellTrace Violet, co-cultured with ρ^0^ astrocytes with and without pyruvate/uridine, and imaged live after 24 hours (Fig 6C). Strikingly, double positive cells with two visually distinct nuclei were observed, consistent with the MStC EtBr-staining showed mtDNA. The proportion of double-positive eGFP+Violet+ cells was approximately 5% of all cells in the co-culture using flow cytometry (Fig 6D), confirming the ability of brain-derived cell types to respond to astrocytes undergoing critical stress by fusing with them.

## Discussion

Since the first observation in 2004^29^, intercellular mitochondrial transfer has been recognized as a true physiological phenomenon that allows both cancerous and non-cancerous cells to take up competent mitochondria in situations where their own mitochondria are failing^26,28,30–35^. There are different ways to accomplish transfer of mitochondria from one cell to another, with tunneling nanotubes, extracellular vesicles and gap junctions most often described. In this paper we show that cell fusion can provide mitochondria to astrocytes. Data presented here show that astrocytes under bioenergetic and biosynthetic stress, can rapidly and permanently reverse severe mitochondrial dysfunction though cell fusion. The fusion events only happened under metabolic stress, and both mesenchymal stromal cells and other astrocytes could be involved.

We used ρ^0^ eGFP astrocytes in a co-culture with stromal cells, as a very simple model of the brain microenvironment. Because of the loss of the mitochondrial genome, ρ^0^ cells require supplementation of the nutrients pyruvate and uridine for survival and division. Removal of pyruvate and uridine from the medium resulted in a bioenergetic crisis specifically to the ρ^0^ eGFP astrocytes in the mixed culture – the CB-MStC cells did not require these supplements. This was an extreme metabolic injury to the ρ^0^ cells – with glycolysis the only source of ATP generation, they must reduce a lot of pyruvate to lactate, in order to re-generate NADH; this reduced pyruvate available for the TCA cycle to produce anabolic intermediaries for production of amino acids and lipids; the lack of electron transport meant dihydroorotate dehydrogenase was inactive and pyrimidine biosynthesis was stopped. An immediate and rapid response was required, which took the form of transplantation of a fully functional mitochondrial electron transport chain from another cell.

The rapid timing of events supports cell fusion as the most likely mechanism for mitochondrial restoration in this model. A ρ^0^ human bone osteocarcinoma cell microinjected with a single mitochondrion was shown to regain a full complement of donor mitochondria in approximately 35 doublings^14^, which would be greater than 70 days for our recipient ρ^0^ eGFP astrocytes, had there been a small number of mitochondria transferred individually. Instead, in this study the full complement of mitochondria was present within 24 hours of onset of bioenergetic injury.

We observed cell fusion between ρ^0^ astrocytes and mesenchymal stromal cells – initially, fused cells contained both nuclei, but over time only a single nucleus was present. We do not know which nucleus ‘won’, the astrocyte or the mesenchymal stromal cell. However, there is significant cross-talk required between the nucleus and mitochondria to coordinate mitochondrial activity. Many hundreds of nuclear-encoded proteins, including those for mtDNA replication and gene expression, such as TFAM, mtSSB and polymerase-γ, are transcribed in the nucleus, translated on cytosolic ribosomes and imported into mitochondria. Signaling between nucleus and non-native mitochondria would need to be well established to allow large scale mitochondrial activity. Some cells expressing both CD105 and cytosolic GFP had 2 nuclei, but in other cells only a single nucleus was evident in the fused cells, and over several days multi-nucleated cells resolved into mono-nucleated. It would be interesting to determine whether the remaining nucleus was CB-MStC derived, whether it was itself a fusion, or whether sufficient signalling could immediately take place between the astrocyte nucleus and the CB-MStC mitochondrial network to allow the astrocyte to retain its own nucleus. The ability of a mitochondria – either as individual organelles or as networks – to communicate with a nucleus is presumably a major determinant in the successful restoration of mitochondrial function.

Cell fusion is not a new concept, and has been demonstrated in neural cells *in vivo*. Research in humans showed that females who received bone marrow transplants from male donors had newly developed Y chromosome-positive neurons.^36^ Similarly, female mice transplanted with GFP+ male bone marrow, developed binucleated GFP+Purkinje cells, and cell fusion was confirmed by fluorescence *in situ* hybridization (FISH) of Y-chromosomes, which were only present in one nuclei^37^. Others found that the incidence of bone marrow/Purkinje cell fusion increased with the inflammatory stimulus lipopolysaccharide (LPS)^38^. This is consistent with reports of bone marrow/muscle fibre fusion, which was observed in response to cardiotoxin-induced muscle injury. In this instance, long-range inflammatory signals were proposed to induce bone marrow derived cells to respond to stress.^39^ Fusion after bone marrow transplant has also been observed in tissues like intestine^40^ and liver^41^, and rapid translation of signals between un-related nuclei and mitochondria may be a critical determinant of efficacy in stem cell transplant and other therapies.

Mitochondrial injury, or loss of mitochondrial function, is observed in many neurodegenerative diseases and other neurological pathologies. To utilize mitochondrial replacement as a therapeutic approach, identifying the best donor cell will be imperative. The bone microenvironment is complex and identifying the precise nature of the mitochondrial donor in this system is difficult. Further, the bone microenvironment may seem irrelevant to the brain microenvironment. However, stromal and perivascular cells in both brain and bone share many of the same markers, including CD105, CD73 and CD90 and are thought to derive from common MSC ancestors.^42^ Adding to the complexity of identifying the donor in this situation, these cells are also likely to be multipotent. The cells characterized loosely here as CB-MStC might be osteoclast or macrophage progenitors, or they might be osteoblasts at various stages of osteogenic differentiation or de-differentiation. Torreggiani *et al*. identified that ∼95% of cells expressing Dmp1, an osteocyte-specific protein were found within bone chip cultures *in vitro*, and many of these cells were highly mobile and/or proliferative. Like the CB-MStC here, the Dmp1+ cells found in the bone chips were CD45^-^ and had a CD105^+^/CD90^+^ subpopulation^43^. In addition to osteogenic lineages, MSCs are precursors to adipogenic, myogenic, and chondrogenic cells. When bone is broken, chondrocytes deposit cartilage matrix as a scaffold for osteoblasts to replace with bone. It has been suggested that chondrocytes can de-differentiate to osteoblasts during this process.^44^ This bidirectional plasticity between terminally differentiated and the multipotent progenitor adds to the complexity in identifying what specific cell types can fuse with ρ^0^ astrocytes, donating themselves and their mitochondria.

The astrocyte stress/injury model was set up to model mitochondrial injury in pathologies like Alzheimer’s, Huntington’s and Parkinson’s diseases, and in astrocyte-derived cancers; to begin to understand the signals and processes that would allow mitochondria to function in a new host cell; and to identify an appropriate donor source of mitochondria to restore function. This system has highlighted the role of cell fusion but does not exclude other mechanisms – in some circumstances, transfer of individual, or small populations of mitochondria may be sufficient to allow restoration of mitochondrial function in astrocytes. However, loss of the entire electron transport chain was a very extreme injury that led to a very extreme response. That response has can allow the dissection of the signals and events that lead to restoration of mitochondrial activity in a new cellular environment.

## Supplementary materials

**Supp Figure 1:**
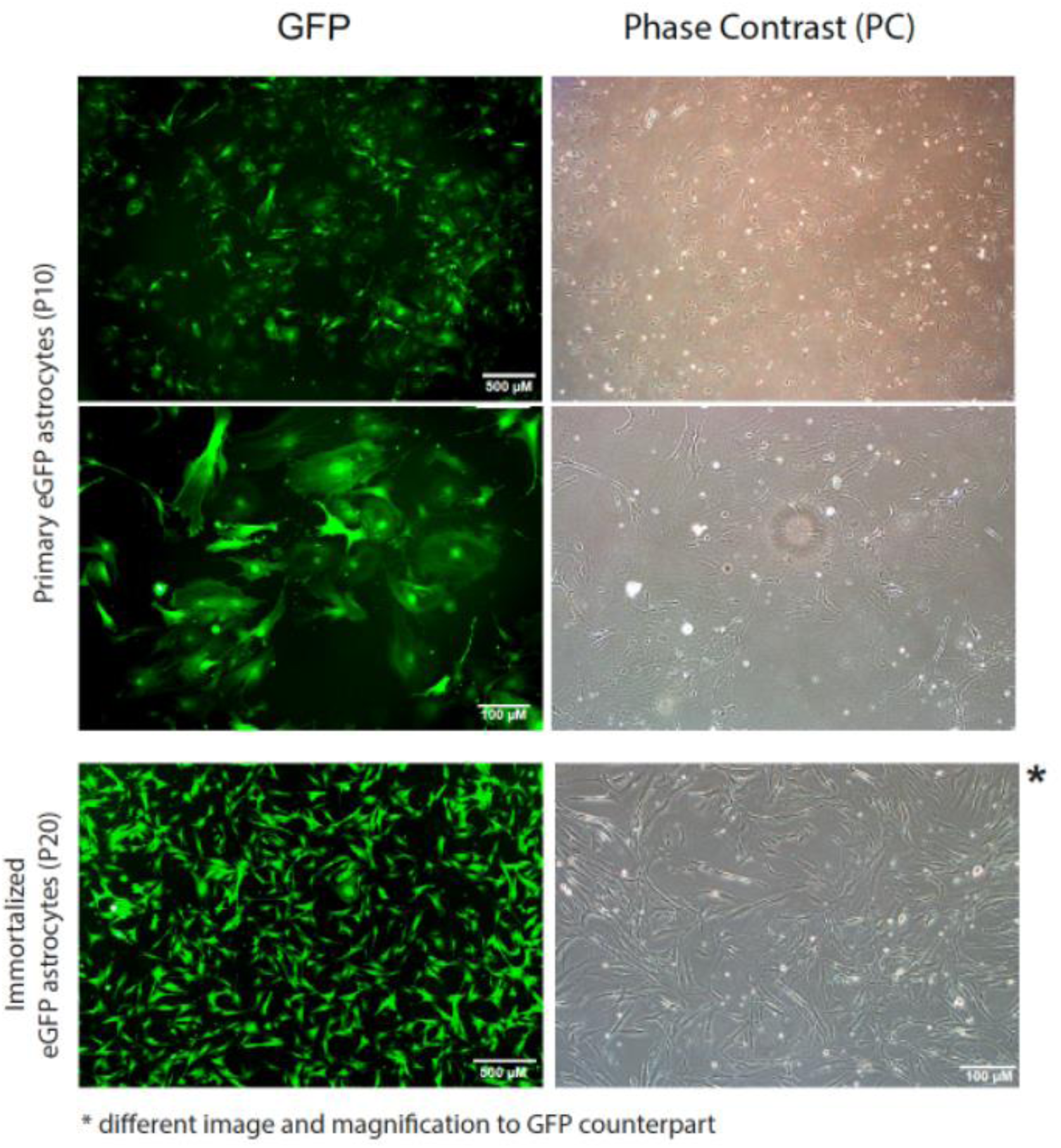
Astrocytes derived from cortical and hippocampal tissue of P0 neonatal eGFP mice. Cultures were passaged multiple times using the 3T3 protocol, which initially selected for the astrocytes away from the neurons in the primary culture. At passage 10 (P10), cells were imaged using phase contrast (right) and fluorescence (left) microscopy. Top panel, scale bar is 500 micron and applies to both fluorescent and phase contrast images. Bottom panel, scale bar is 100 micron, and applies to both fluorescent and phase contrast images. Cells continued to be passaged and spontaneously immortalised approximately passage 18. At passage 20 (P20) cells were imaged again. Fluorescent image scale bar, 500 microns. Phase contrast image scale bar, 100 microns.

**Supp Fig 2:**
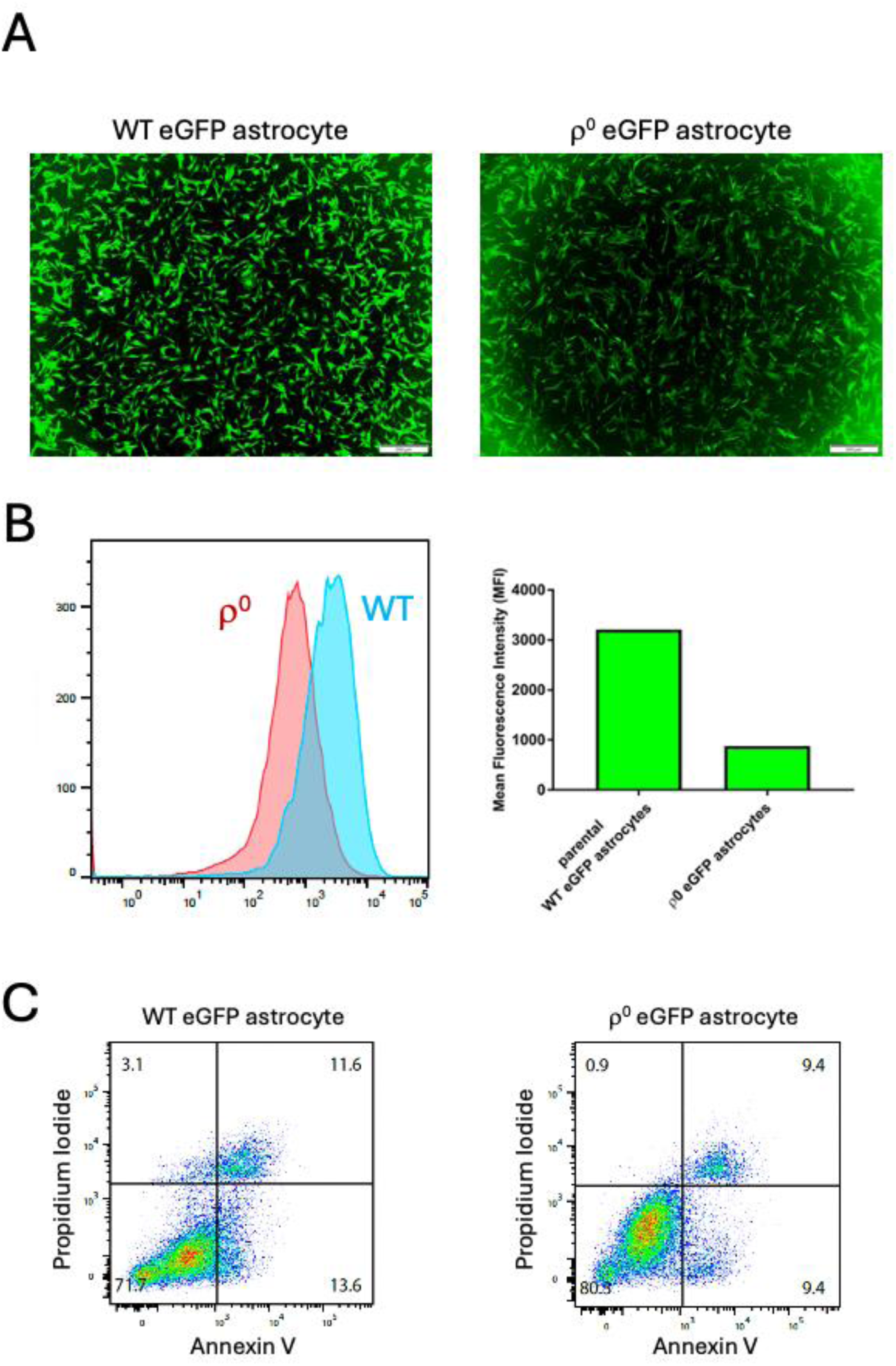
GFP expression and basal apoptosis of wildtype and mitochondrial DNA deficient r0 cells. (A) eGFP expression was imaged by fluorescence microscopy in both cell types. (B) eGFP fluorescence was quantified by flow cytometry in live cells, and the median fluorescence intensity of the r0 cells was approximately 1/3 of the wildtype cells. (C) Apoptosis was measured using annexin V staining with propidium iodide (PI), with similar proportions of AnnexinV+PI-cells (early apoptosis, lower right quadrant) and AnnexinV+PI+ cells (late apoptosis, upper right quadrant).

**Supp Fig 3:**
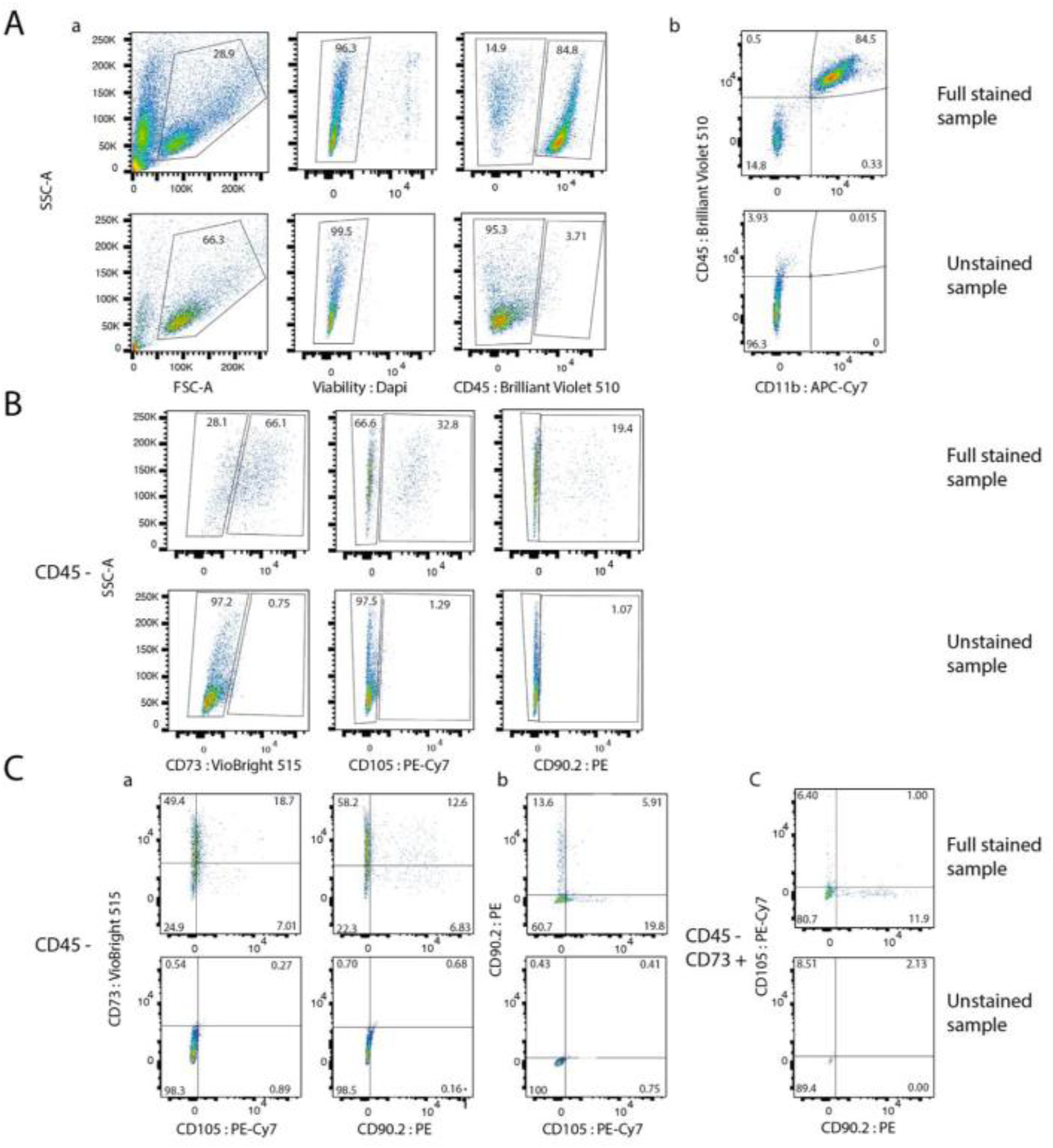
The full staining and gating panel for compact bone derived mesenchymal stromal cells (CB-MStC). (A) Gating strategy for (a) CD45+ and CD45-cells, and (b) CD45+ CD11b+ cells. The unstained controls for gating shown in the lower panels. (B) Single staining for mesenchymal stem cell markers CD73, CD105 and CD90.2 on CD45- cells, with unstained controls in the lower panels. (C) Double staining of CD45- cells for (a) CD73 versus CD105 and CD90.2; and (b) CD90.2 versus CD105. (c) CD45-CD73+ cells double-stained for CD90.2 versus CD105. Unstained samples in the lower panel.

---

**Supplementary Table 1.**
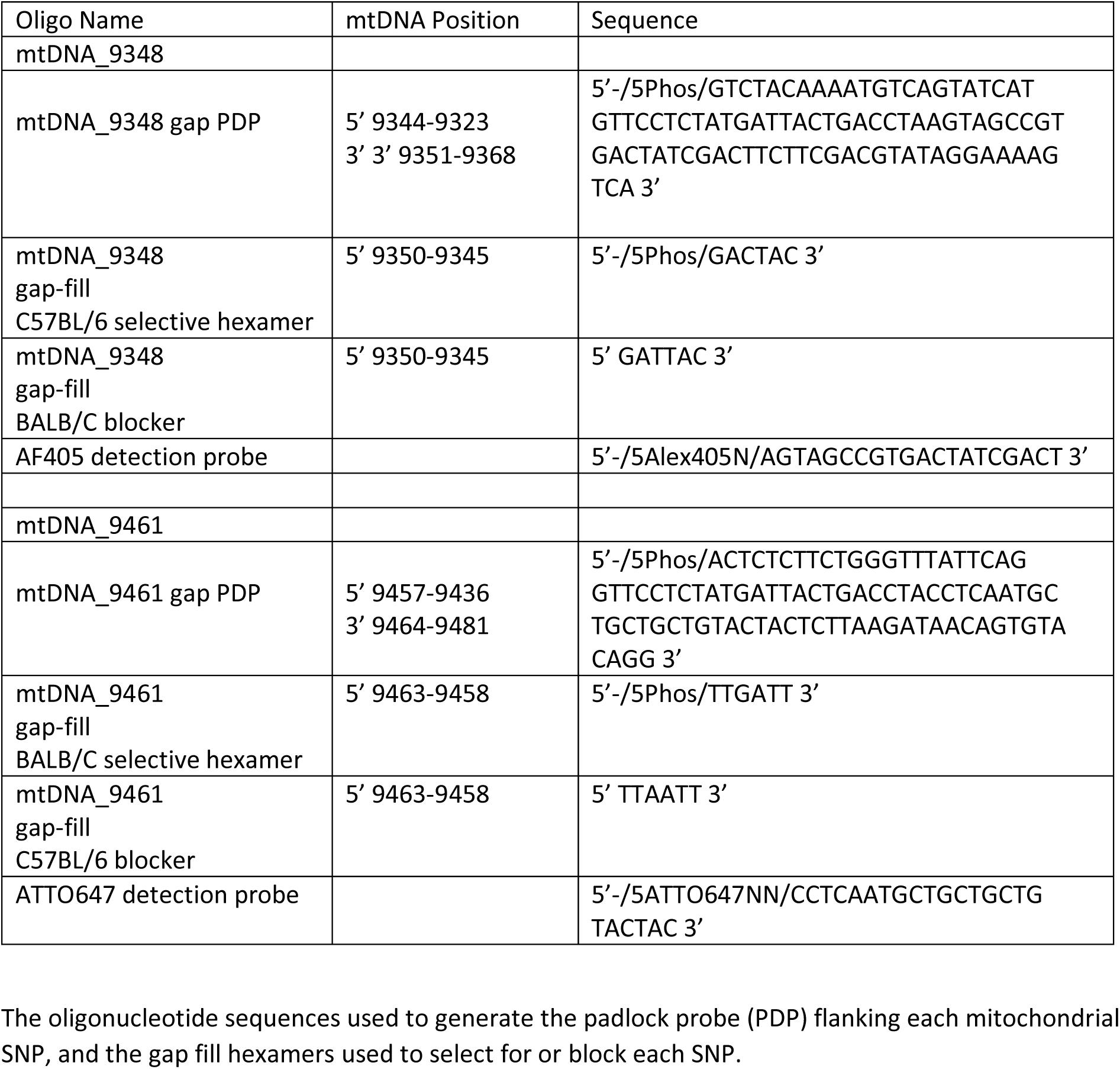
Oligonucleotides for *in situ* target-primed RCA of mtDNA_9348 and 9461 by gap-fill ligation.

